# Hyperglycemia transcriptionally regulates the paranodal protein (Caspr1) in retinal neurons and modulates neurite extension

**DOI:** 10.64898/2026.01.03.691327

**Authors:** S Sharma, S Shree, G Suresh, H Chakravarthy, M Schmelz, KK Bali, Sushma Dagar, BS Sahu, I Kaur, V Devanathan

## Abstract

Hyperglycemia is a hallmark of diabetes, affecting neuronal structure and function by altering molecular signalling pathways. Here, we explore the role of hyperglycemia in regulating Caspr1 expression and its downstream effects on neurite outgrowth. Caspr1, a critical protein implicated in neurodegenerative diseases, was found to be significantly downregulated in N2a and 661W cell lines cultured under hyperglycemic conditions (25mM glucose) and, as a result, promoted neurite outgrowth. Knockout of Caspr1 using CRISPR-Cas9 further confirmed its inhibitory role on neurite outgrowth, as Caspr1-deficient cells exhibited enhanced neurite elongation. Caspr1 downregulation was mediated by decreased expression of C/EBPα, a key transcription factor with a binding site on the Caspr1 promoter. Overexpression of C/EBPα restored Caspr1 promoter activity and mRNA levels, establishing C/EBPα as a critical regulator. Additionally, hyperglycemia was observed to inhibit Akt phosphorylation, which further contributed to Caspr1 downregulation. Adding insulin to the culture medium under hyperglycemic conditions shows inhibition of Akt phosphorylation and downregulation of Caspr1, resulting in a shorter length of neurites in retinal neurons. *In vivo,* studies in diabetic mouse models and diabetic patient samples demonstrated reduced expression of Caspr1 in retinal tissues. These results suggest that hyperglycemia regulates Caspr1 expression through Akt and C/EBPα pathways, promoting neurite outgrowth in retinal neurons. In contrast, adding insulin to the medium under hyperglycemia downregulates the Caspr1 expression and reduces neurite length in retinal neurons. Targeting this pathway may offer new therapeutic approaches to mitigate neurodegeneration in diabetic retinopathy.

## Introduction

In the central nervous system, neuritogenesis is an intricate process that incorporates multiple signals to generate transcriptional and morphological changes. During this process, immature neurites develop initially, which is known as neurite outgrowth, and these neurites continue to grow into a single axon and many dendrites. Neurite outgrowth is meticulously controlled because it is crucial for the proper development of the organism (Bromberg et al., 2008; Huang and Reichardt, 2001). Several neurodegenerative diseases of the central nervous system (CNS), such as Alzheimer’s disease, temporal lobe epilepsy, and retinal degeneration, stimulate adult-onset neuritogenesis (Scheibel and Tomiyasu, 1978; Jones et al., 2003; Marc and Jones, 2003; Elliott et al., 2003; Dadon-Nachum et al., 2011). One of the areas of focus of the research on the preventive and therapeutic impacts of neurodegenerative illnesses is neuritogenic activity. By encouraging the formation of neurites from neuronal cells, neuritogenic chemicals hold the promise of therapeutic efficacy in treating neuronal damage (More et al., 2012; Ling-Sing Seow et al., 2013). Neuritogenesis in the adult CNS may be corruptive and undermine vision rescue techniques in retinal degeneration, including age-related macular degeneration and retinitis pigmentosa (Marc et al., 2003, 2007; Lin et al., 2012). Understanding the initiators of aberrant neuritogenesis and discovering molecular targets that govern corruptive network formation is crucial for optimising therapy outcomes for retinal degeneration and CNS disorders.

Contactin-associated protein 1 (Caspr1), also referred to as paranodin, is a cell adhesion molecule and is essential for the maintenance of the node of Ranvier in myelinated neurons. It forms a complex with Contactin and Neurofascin-155, and this interaction helps establish and maintain the precise positioning of the myelin sheath relative to the axon, ensuring proper propagation of action potentials along the axon (Tang et al., 2020). Caspr1 acts as a negative regulator of neurite outgrowth by interacting with prion protein (Devanathan et al., 2010). Caspr1 is known to participate in neurodegenerative diseases like Alzheimer’s disease and multiple sclerosis. Some studies show that Caspr1 may form a complex with γ-secretase, indicating that it is involved in the pathogenesis of Alzheimer’s disease (Hur et al., 2012). Others have demonstrated that Caspr1 is localised around amyloid plaques in the mouse cerebral cortex and interacts with the amyloid precursor protein. Caspr1 overexpression reduces the production of Amyloid-β (Aβ) 40 and Aβ42 (Fan et al., 2013). In multiple sclerosis, morphological disorganization has been observed in nodal, paranodal and juxta-paranodal regions in myelinated neurons. During the initial phases, Caspr1 and NF-155 clusters at the paranode region are observed to be elongated, while voltage-gated K^+^ channels are dispersed and gradually start to overlap with the paranodal area (Coman et al., 2006). It has also been observed that demyelination leads to a decrease in Caspr1 expression in lesioned areas, while with the formation of a new myelin sheath, Caspr1 reappears. Changes in Caspr1 expression may foreshadow abnormalities with saltatory conduction and further myelin loss (Coman et al., 2006; Howell et al., 2006; Zou et al., 2017). Caspr1 is well-studied in the brain for its function in forming and maintaining myelin sheath, organization of nodes and paranode region. Caspr1 is also present in the retina, which has non-myelinated neurons, although its role and regulation remain unclear. Our earlier research demonstrated that hyperglycemia significantly promotes neurite growth in cultured retinal neurons while suppressing the expression of neuronal transmembrane proteins such as Caspr1 and Contactin1 (Sharma et al., 2019).

In this study, for the first time, we have shown that Caspr1 is transcriptionally regulated under hyperglycemic conditions in retinal neurons and is a target gene of a glucose-sensitive transcription factor, CCAAT/enhancer binding protein (C/EBP) α. Furthermore, we observed that hyperglycemia downregulates the conversion of Akt to phospho-Akt as well as C/EBPα expression; as a result, we observed longer neurite outgrowth in retinal neurons. We also observed neurite length extension in Caspr1 knockout cells, which shows that Caspr1 negatively regulate neurites. Subsequently, we found that in type-1 diabetic mice and high-fat diet mice (HFD)-induced obese/ hyperglycemic, the retina has significantly reduced expression of Caspr1 and C/EBPα compared to the control mice. Likewise, Caspr1 expression was downregulated in type II diabetic patients. Combined, our studies firmly establish a novel regulatory mechanism for neurite extension by Caspr1 in hyperglycemic conditions. Targeting this pathway could help develop new therapeutic strategies to prevent neurodegeneration in the diabetic retina.

## Materials and methods

### Cell culture and other reagents

The Neuro-2a (N2a) cell line was obtained from ATCC, and the 661W cell line was a kind gift from Dr. Ghanshyam Swarup, CCMB, Hyderabad, India (Sayyad et al., 2017). The cell lines were cultured in DMEM supplemented with 10% fetal bovine serum (FBS) (Thermo Scientific) and 1X antibiotic solution (Penicillin/Streptomycin, Sigma). Cells were cultured in 5% CO_2_ incubator at 37°C, in 5mM (normoglycemia) and 25mM (hyperglycemia) glucose-containing media for 48 hours. For inhibition of gene transcription, cells were grown in 5mM and 25mM glucose-containing DMEM media for 48 hours. Cells were treated with Actinomycin D (10ug/ml, Sigma) and incubated for another 6 hours. For inhibition of Akt, cells were incubated with 10-NCP (Sigma) (AKT inhibitor) for 24 and 48 hours.

### Primary retinal culture

C57BL6/j mice were sacrificed using isoflurane. Eyeballs were carefully removed and kept on ice in sterile HBSS (Gibco). Under the dissection microscope, the eyeball was punctured, and the retina was extracted from the eyecup in HBSS solution. Retinal tissue was washed and digested with 0.25% trypsin (Gibco) for 10 minutes at 37°C. After complete digestion, DMEM (Gibco) complete media containing 10% FBS (Gibco) and 1% Penicillin/Streptomycin (Sigma) was added to terminate the digestion reaction. The cell suspension was filtered through a 40μm cell strainer (Himedia) and centrifuged at 1000 rpm for 5 minutes. The cell pellet was resuspended in DMEM complete media. Cells were counted by using trypan blue (Gibco). After counting, cells were cultured in 5mM glucose-containing DMEM (normoglycemia) and 25mM glucose-containing DMEM (hyperglycemia) for 10 days. Half medium was changed every alternate day. Cells were used on day 10 for further experiments.

### Quantitative real-time PCR (qPCR)

To perform RT-PCR, total RNA was isolated from cells using TRIzol (Invitrogen). 1μg RNA was converted to cDNA using an iScript cDNA synthesis kit (Bio-Rad). 2μl of cDNA was used to perform qPCR to check the expression of specific genes using iTaq Universal SYBR Green supermix (Bio-Rad). Results were normalised to β-actin. The primers used are given below.

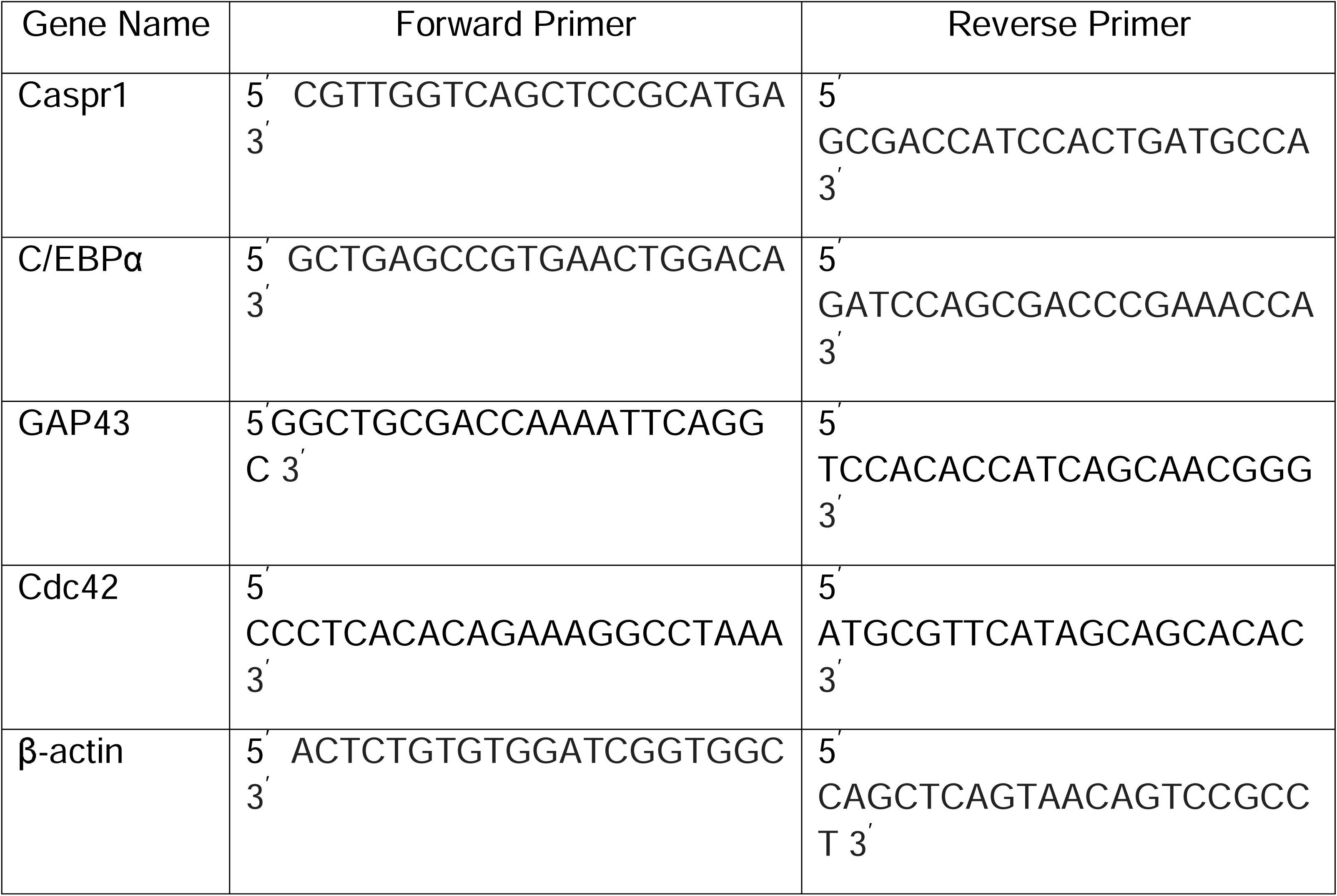

### Plasmid construction

The mouse Caspr-1 promoter sequence was extracted from UCSC ( https://genome.ucsc.edu/ ), and primers were designed to clone the promoter sequence:

Forward- 5^’^ AAACCCGGGCCCAGAGGGAGCGCGAAGCCG 3^’^

Reverse- 5^’^ GGGCTCGAGCCTCCCTCTCCCTCCCTCTCTCTGC 3^’^

The Mouse Caspr-1 promoter was PCR amplified using genomic DNA isolated from N2a cells. The PCR amplified promoter region was cloned into the pGL2 basic vector. C/EBPα coding sequence (CDS) was amplified from N2a cDNA, which was synthesised using gene-specific primers using a cDNA synthesis kit (BioRad).

C/EBPα specific primer sequences are given below:

Forward- 5^’^ GACGAATTCATGGAGTCGGCCGACTTCTACGAGG 3^’^

Reverse- 5^’^ GAATCTAGATCACGCGCAGTTGCCCATGGCCTTG 3^’^

Amplified C/EBPα was cloned into the pcDNA3.1/HisC plasmid and used for further experiments.

### Reporter assay

Caspr-1 luciferase assay was performed per the manufacturer’s instructions (Promega, Madison, WI, USA). N2a and 661W cells were lysed using 1X reporter lysis buffer after 48 hrs of transfection, then centrifuged at 12,000 rpm, 4°C for 5 minutes, and the supernatant was collected. Caspr1 promoter luciferase assay was performed using Luciferase Assay System (Promega) by adding 20μl of cell lysate to 100μl of luciferase assay reagent and measuring luminescence using a luminometer. Results were normalised against an internal control, β-galactosidase activity. The β-galactosidase activity was measured using the β-galactosidase Enzyme Assay System (Promega) at 420nm.

### Western blot and antibodies used

Cells were lysed in RIPA lysis buffer. Samples were incubated on ice for 30 minutes and centrifuged at 15,000 g. After centrifugation, the supernatant was extracted, and protein concentration was determined using a BCA kit (Thermo Fisher). Samples were mixed with sample buffer and boiled for 5-10 minutes at 95°C. Proteins were separated by electrophoresis on 8, 10, or 12% SDS-polyacrylamide gels and transferred to a nitrocellulose membrane. Membranes were blocked with 3% BSA in TBS, incubated with primary antibodies overnight at 4°C with shaking and washed with TBST. Primary antibodies were detected with corresponding HRP-conjugated secondary antibodies applied for 1 hour in 3% BSA in TBS. After washing, secondary antibodies were visualised with chemiluminescence. Antibodies against Caspr-1 were procured from Sigma and Abcam, while those against AKT, p-AKT, and C/EBPα were procured from Cell Signalling Technologies. α-Tubulin and β-actin were purchased from Abcam.

### Caspr-1 knockout cell line

Caspr-1 gene sequence was submitted to crispr.mit.edu for the design of sgRNA (short guide RNA which directs the Cas9 nuclease to a cleavage site in the genome). Two sgRNAs with high scores were chosen for the target, showing the lowest scores for mismatches. The sgRNAs sequences were as follows:

sgRNA 1: 5^’^ CACCGCCACTGGCCTGGAACCCACG 3^’^

sgRNA 2: 5^’^ CACCGTGTGACGCTTGAACTGCAAG 3^’^

These sgRNAs targeted exons 4 and 5 of the Caspr-1 gene. These sgRNAs were cloned into BbsI-digested pU6-(BbsI) _CBh-Cas9-T2A-mCherry vector (Addgene), and vectors were transformed into competent *E.coli* cells. Clones were confirmed by colony PCR by using specific primers.

**For Oligo1**

Forward primer: 5^’^ CACCGCCACTGGCCTGGAACCCACG 3^’^

Reverse primer: 5^’^ GGGGGCGTACTTGGCATAT 3^’^

**For Oligo2**

Forward primer: 5^’^ CACCGTGTGACGCTTGAACTGCAAG 3^’^

Reverse primer: 5^’^ GGGGGCGTACTTGGCATAT 3^’^

After confirmation of clones containing both sgRNAs, plasmids were isolated from each clone, and Gibson assembly reaction was performed to obtain both sgRNAs into a single vector containing Cas9. Gibson assembly reaction was performed using Gibson master mix (detailed composition of master mix in Appendix of thesis), which contains T5 exonuclease (NEB) and Phusion DNA polymerase (NEB). The reaction was incubated at 50°C for 1 hour. After Gibson assembly, plasmids were transformed into competent *E.coli* cells. Clones were confirmed by colony PCR using specific primers:

Forward primer: 5^’^ CACCGCCACTGGCCTGGAACCCACG 3^’^

Reverse primer: 5^’^ AAACCTTGCAGTTCAAGCGTCACAC 3^’^

In addition to confirmation by colony PCR, the clones were further confirmed using Sanger sequencing. The positive clone was transfected using Lipofectamine 3000 (Thermo Scientific) into the N2a cell line, passage number 16, and checked for mCherry expression. The transfected cells were seeded in a 96-well plate, diluted so that each well contained a single cell, and clones were allowed to grow for two weeks. Next, genomic DNA was isolated from each well, and screening was done for knockout cells by PCR using Caspr-1 gene-specific primers between exon 4 and exon 5:

Forward primer: 5^’^ CCTTCACTACCACTTTACGGCT 3^’^

Reverse primer: 5^’^ AATGGGAAAGAGCAACCTGAGT 3^’^

Expression of Caspr-1 was also checked by Western blot using Caspr-1 antibodies (Abcam) 1:250 dilution to confirm Caspr-1 knockout in N2a cells.

### Neurite outgrowth assay

Cells were cultured and fixed with 4% formaldehyde. After fixing, the cells were subjected to trypan blue staining for 15 minutes at room temperature. To quantify neurite length, cells were visualised under 10X magnification, and the total length of all neurites was determined by the NeuronJ software plugin in ImageJ software. Neurite length measurements from at least 100 cells per dish were recorded from randomly chosen fields. Each experiment was repeated three times.

### Immunocytochemistry

Immunofluorescence staining was performed as described in (Sharma et al., 2019). The cells were seeded on top of the coverslip inside the six-well plate and grown in DMEM media. After attaining 80% confluency, cells were fixed with 4% paraformaldehyde for 10–15 minutes and then washed with ice-cold PBS. Post-fixing, the cells were permeabilised with 0.2% Triton X-100 for 10 min and then washed thrice with ice-cold PBS. The cells were then blocked with 1% BSA in Phosphate Buffered Saline with Tween-20 (PBST) for 1 hour at room temperature. After 1 hour, the cells were treated with primary antibodies (anti-Caspr1 rabbit antibody, Sigma, and anti-β-III tubulin rabbit antibody, CST) prepared in 1% BSA in PBST and incubated overnight at 4°C. After incubation, cells were washed thrice with ice-cold PBS for 5 minutes. The cells were then incubated with secondary antibodies prepared in 1% BSA in PBST (Alexa Fluor-488 anti-mouse antibody and Alexa Fluor-594 anti-rabbit antibody, Invitrogen) at room temperature for 1 hour in the dark. The cells were washed thrice with ice-cold PBS for 5 minutes each in the dark. The coverslips with processed cells were mounted onto a slide and observed under an IX83 inverted epifluorescence microscope (Olympus).

### Type-1 Diabetic mouse model

Type I diabetic mice were a kind gift from Dr. Nitin Agarwal, Heidelberg University, Germany. Type I diabetes was induced as described in (Agarwal et al., 2020). In 8-week-old C57BL6/j mice, diabetes was induced by 5 sequential days of intraperitoneal (i.p.) injections of Streptozotocin (STZ) at a low dose (60 mg/kg/d) in citrate buffer. In the untreated group, only citrate buffer was injected. Throughout the duration of the study, blood glucose levels were checked once per week in the afternoon using a glucometer (Accu-Chek, Roche Diagnostics). Mice with blood glucose levels higher than 350 mg/dl were considered diabetic mice. To achieve uniformity and consistency, the blood glucose levels were kept within a range of 350 and 500 mg/dl throughout the duration of the experiment by subcutaneous injection of insulin.

### High-fat diet mouse model

The high-fat diet mice were provided by Dr. Bhavani Shankar Sahu, NBRC Manesar, India. All the animal experiments were performed after approval by the National Brain Research Centre (NBRC) animal ethics committee. Six-week-old C57BL/6J mice (n=25 males) were group-housed after weaning (siblings in one cage) and kept on a 12-hour light-dark cycle in the animal house facility of NBRC.

### Type II diabetic patient

Type II diabetic patients’ samples were provided by Dr. Inderjeet Kaur, Kallam Anji Reddy Molecular Genetics Laboratory, Brien Holden Eye Research Centre, L V Prasad Eye Institute, Hyderabad, India. All the experiments were performed after approval by L V Prasad Eye Institute, Hyderabad, India.

### Statistical Analysis

Data are presented as mean ± SEM. Results were analysed for statistical significance by Student’s *t*-test (Excel). The degree of significance is depicted as follows: \**P*<0.05, \*\**P*< 0.01, and \*\*\**P*<0.001.

## Results

### Hyperglycaemia downregulates Caspr-1 expression

To examine the effect of glucose on the expression of Caspr-1, N2a and 661W cell lines were cultured in two different concentrations of glucose-containing media (5 mM and 25 mM). It was observed that high glucose concentration in the culture medium significantly downregulated Caspr1 mRNA expression in both cell lines (Figure 1A). Downregulation in the expression of Caspr1 protein in western blot and Caspr1-immunostained cells was observed under hyperglycemic conditions in N2a and 661W cells (Figure 1C, D). Further, the effect of glucose on cultured primary retinal cells was studied, and similar results were observed (Figure 1B). Treatment with Actinomycin D (10μg/ml) for 5-6 hours, a transcriptional inhibitor, effectively blocked the expression of Caspr1 mRNA and the mRNA level was significantly reduced in 5mM and 25mM glucose medium in N2a and 661w cells (Figure 1E). These observations suggest transcriptional dependency of Caspr1 expression in N2a and 661W wells.

**Figure 1.**
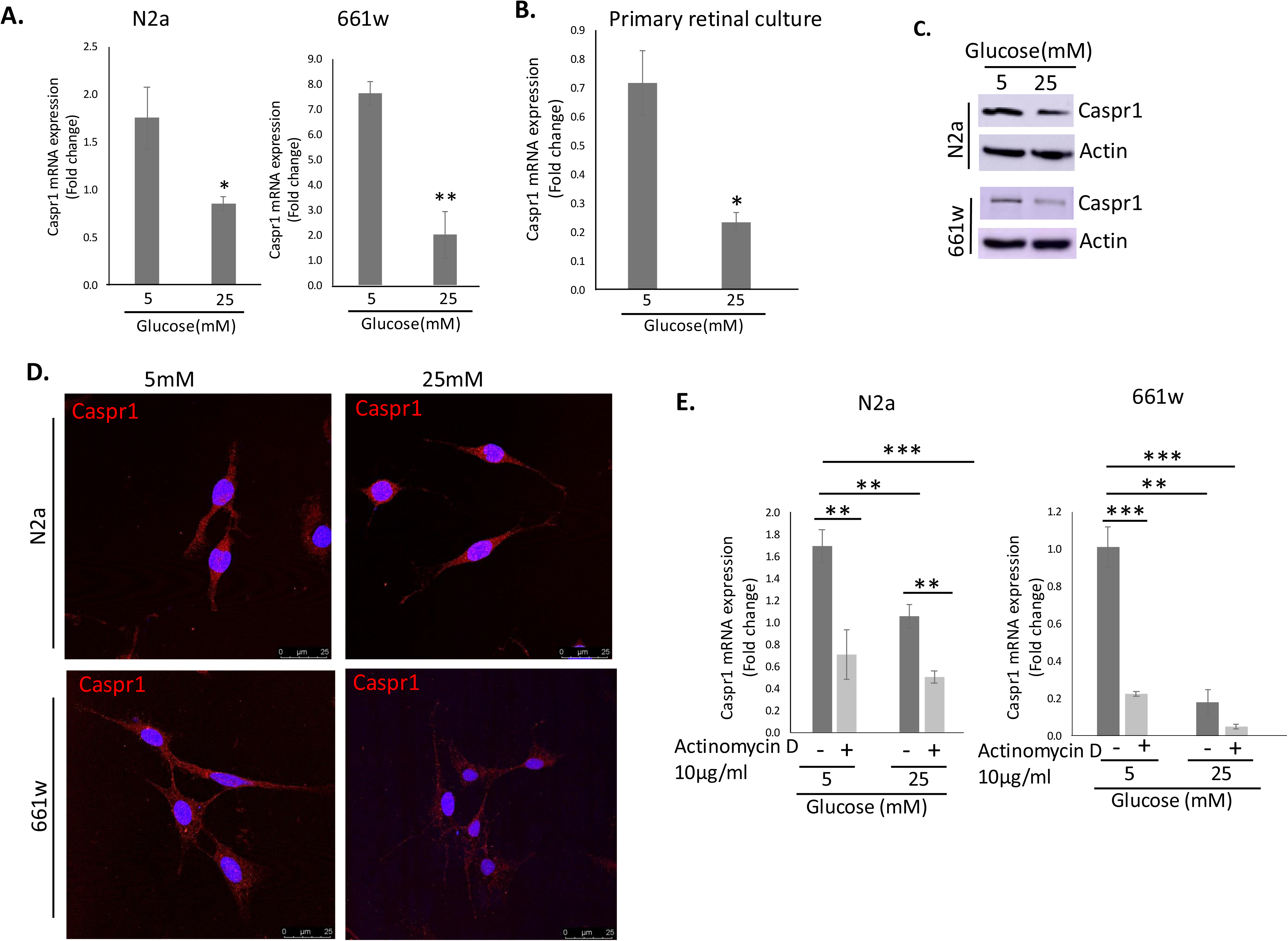
Hyperglycaemia downregulates Caspr1 expression. (A) qPCR analysis for Caspr1 mRNA expression in N2a and 661w cells grown in 5mM and 25mM glucose-containing media. (B) qPCR analysis for Caspr1 mRNA expression in cultured retinal cells grown in 5mM and 25mM glucose-containing media. (C) Western blot analysis for the Caspr1 protein expression in N2a and 661w cells grown in 5mM and 25mM glucose-containing media. (D) Expression of Caspr1, red in N2a and 661w cells in 5mM and 25mM glucose by immunofluorescence. (E) Effect of actinomycin D on the mRNA expression of Caspr1 in N2a and 661w cells in medium containing 5mM and 25mM of glucose. N=4. *P<0.05, **P<0.01, ***P<0.001.

### Hyperglycemia promotes neurite outgrowth

To understand the structural changes induced by altered glycemic conditions in neurons, neurite length of neurons grown in normal and hyperglycemia conditions were measured. In short, cells were grown in different glucose conditions for 48 hours, and a neurite outgrowth assay was performed. It was observed that cells had significantly longer neurites under hyperglycemic conditions than in normoglycemic conditions; N2a cells (5mM: 29.4 ± 0.47 μm, 25mM: 47.7 ± 0.97 μm (Figure 2A, B) and 661W cells (5mM 16.8 ± 0.15 μm, 25mM 49.6 ± 0.37 μm) (Figure 2D, E). Further, when the data were segregated depending on the neurite length of cells, it was observed that in normoglycemia, most cells had shorter neurite lengths (20-40μm), while in hyperglycemia, cells had longer neurites (60-80μm) in both cell lines (Figure 2C, F). Since hyperglycemic conditions (25 mM glucose) promote neurite outgrowth, we quantified the expression of Gap43 and Cdc42, which are known to promote neurite outgrowth (Govek et al., 2005). The results demonstrate that hyperglycemia upregulates the expression of these proteins to promote neurite length in both cell lines (Figure 2G).

**Figure 2.**
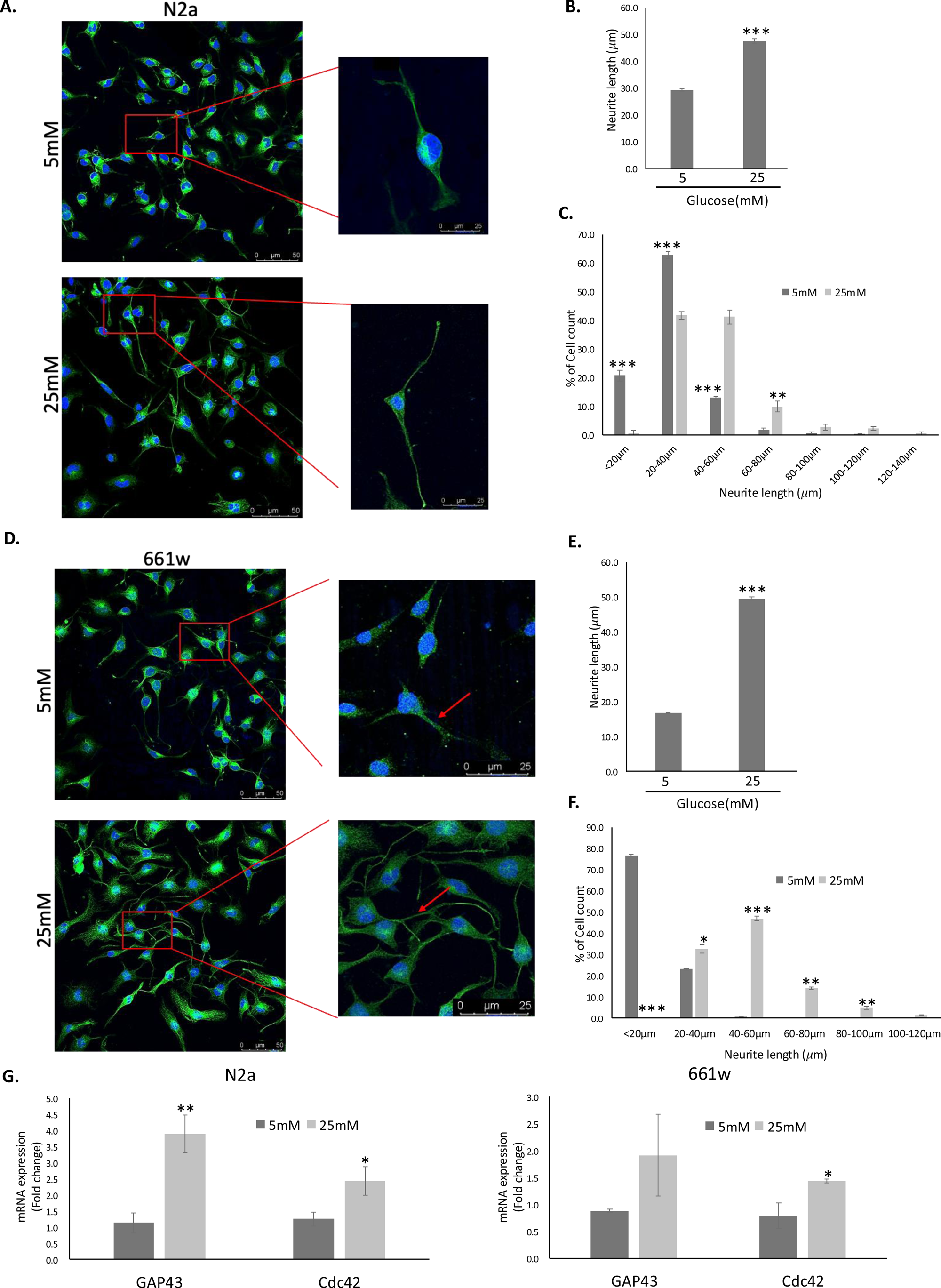
Hyperglycemia significantly promotes neurite outgrowth. (A) and (D) Representative 40X *β*III tubulin (green) immunostained images showing the length of neurite (zoomed image, red arrow) in N2a and 661W cells cultured in 5mM and 25mM glucose medium, respectively. (B) and (E) Quantification of total neurite length shows a significant increase in 25mM glucose compared to 5mM glucose-containing media in N2a and 661w cells. (C) and (F) The percentage of cell count shows that more cells have longer neurites (40-80*μ*m in N2a and 40-100*μ*m in 661w cells) in 25mM than the 5mM glucose medium in N2a and 661w cells. N=6. (G) qPCR analysis of GAP43 and Cdc42 mRNA expression in N2a and 661w cells grown in 5mM and 25mM glucose medium. N=3. *P<0.05, **P<0.01, ***P<0.001.

### Caspr1 inhibits neurite outgrowth

To investigate if Caspr1 expression has an influence on the neurite outgrowth, Caspr1 knockout N2a cells were created using the CRISPR-Cas9 twin-guide system (Supplementary Figure 1). Caspr1 knockout in cells was confirmed by checking for the deleted region in the genome of cells by PCR and by examining the loss of Caspr1 expression at the protein level in knockout cells versus wild-type cells by Western Blot (Figure 3A, B). Microscopic and morphological analyses revealed that ΔCaspr1^+/-^ cells had significantly longer neurites that Caspr1^+/+^ wild-type N2a cells (Figure 3C and 3D). To confirm this observation, neurite lengths in wild-type and knockout cells were quantified and compared using NeuronJ software. ΔCaspr1^+/-^cells had significantly longer neurites than Caspr1^+/+^ wild-type cells (Figure 3D). Additionally, the live cell imaging of wild-type and knockout cells also showed that knockout cells grow neurites much faster and longer than wild-type cells (Video 1). The above results indicate that Caspr1 is a negative regulator of neurite outgrowth.

**Figure 3.**
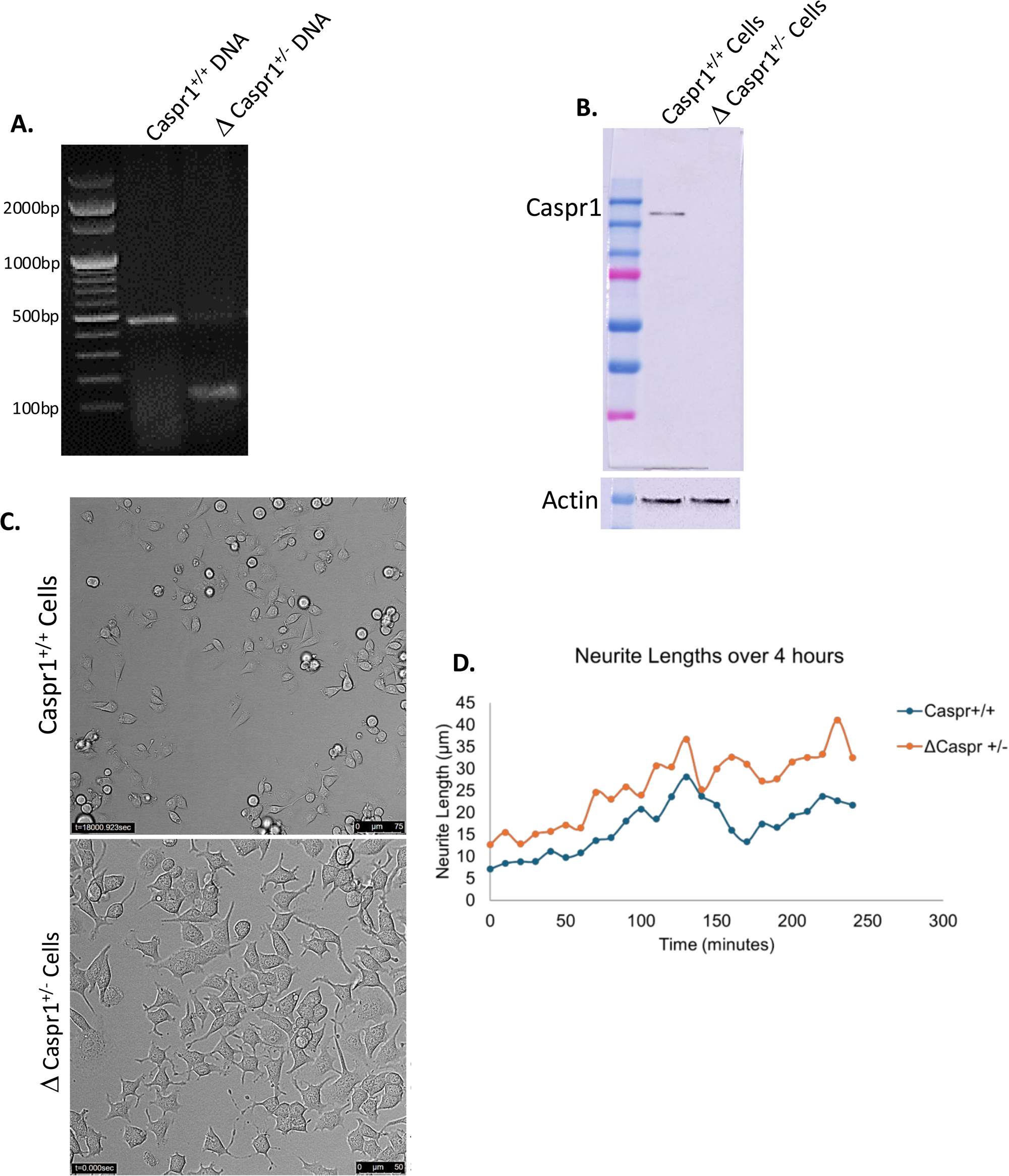
Caspr1 mutation promotes neurite elongation. (A) Screening of the Caspr1 knockout cell line using PCR. (B) Western blot analysis for Caspr1 protein expression in Caspr1 knockout cells. (C) Representative bright field images showing the morphology and neurite length in wild-type and knockout cells. (D) Quantification of total neurite length shows a significant increase in Caspr1 knockout cells over 4 hours.

### Hyperglycaemia downregulates Caspr1 promoter activity

To investigate whether the decrease in Caspr1 mRNA by glucose was due to a decrease in its promoter activity, the mouse Caspr1 putative promoter (-1 to -2000 bp, full length) was cloned in pGL2 basic vector (Supplementary Figure 2). The inducibility of the Caspr1 promoter was examined in N2a and 661W cells grown under different glucose conditions (5mM and 25mM) after transient transfection of the mouse Caspr1 promoter reporter plasmid and by measuring luciferase activity. Caspr1 promoter luciferase activity was significantly higher in cells cultured in 5mM glucose-containing media than in 25mM glucose (Figure 4A). Consistent with these results, we also noticed a similar pattern with the Caspr1 protein expression (Figure 1C).

**Figure 4.**
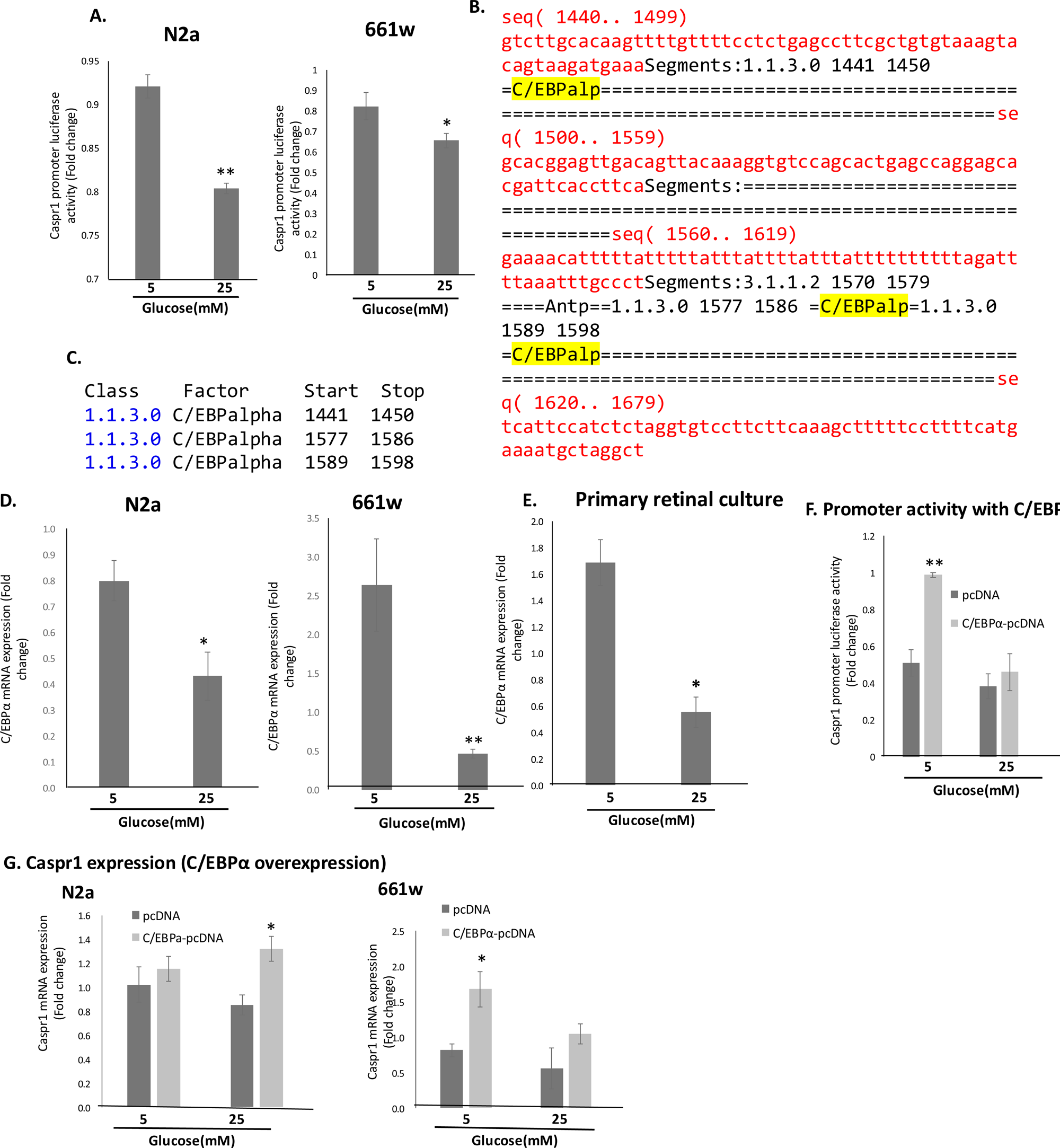
C/EBP*α* regulates the transcription of Caspr1. (A) Caspr1 Promoter (full length) luciferase activity in N2a and 661w cells grown in a medium containing 5 and 25 mM glucose. (B, C) The binding site of C/EBP*α* to the promoter of the Caspr1 gene. (D, E) qPCR analysis of C/EBP*α* mRNA in N2a, 661w and cultured retinal cells grown in 5mm and 25mM glucose-containing medium. (F) Caspr1 promoter luciferase activity in N2a cells transfected with the pcDNA or pcDNA-C/EBP*α* expression plasmid and cells were grown in 5mM and 25mM glucose. (G) qPCR analysis of Caspr1 mRNA expression in N2a and 661w cells transfected with pcDNA or pcDNA-C/EBP*α* in the presence of 5mM and 25mM glucose-containing medium. N=3. *P<0.05, **P<0.01.

### C/EBP**α** regulates the transcription of Caspr1

The effect of glucose on the expression of specific transcription factors was examined. Using the Alibaba2 program (http://gene-regulation.com/pub/programs/alibaba2/), the binding sites of transcription factors on the Caspr1 promoter were predicted. After analysing the results, CCAAT/enhancer binding protein (C/EBP) was selected over other transcription factors because it is a glucose homeostasis-sensitive transcription factor. Results showed that C/EBPα has a binding site at the mouse Caspr1 promoter between -1400 and -1600 bp (Figure 4 B, C). The effect of hyperglycemia on C/EBPα was studied, and RT-PCR data showed significant downregulation of C/EBPα expression at the mRNA level (Figure 4D, E). Scanning of mouse Caspr1 putative promoter sequence (-1 to -2000bp) revealed three potential binding sites for C/EBPα (-1441, -1577, and -1589) (Figure 4C). To elucidate whether Caspr1 is directly regulated by C/EBPα, Caspr1 promoter luciferase activity was performed in N2a and 661W cells co-transfected with C/EBPα expression plasmid (Supplementary Figure 3) and Caspr1 promoter reporter construct. C/EBPα overexpression readily stimulated Caspr1 promoter luciferase activity over the baseline, even in the presence of high glucose (Figure 4F). We also observed upregulation in Caspr1 mRNA expression in N2a and 661W cells overexpressing C/EBPα (Figure 4G). In summary, C/EBPα forms a complex with the Caspr1 promoter and drives the transcription of Caspr1, which is affected in hyperglycemic conditions.

### Hyperglycemia affects the Akt-dependent expression of Caspr-1

The PI3K-Akt pathway, activated by neurotrophins like insulin, has been recognised as a key signalling pathway for neuronal survival. The impairment of Akt-mediated signal transduction plays a significant role in the development of various neurodegenerative diseases, including Parkinson’s disease, Alzheimer’s disease, Huntington’s disease, and others (Rai et al., 2019; Read and Gorman, 2009). Hyperglycemia alters the activation of Akt and affects endothelial cell proliferation and survival (Varma et al., 2005). To check the effect of hyperglycemia on Akt in neuronal cells, cells were grown in different glucose conditions, and the expression of signalling molecules such as Akt and pAkt was examined. High glucose concentration was observed to downregulate the expression of pAkt at the protein level in both cell lines (Figure 5A, B). Next, we examined whether AKT is involved in Caspr1 expression; When cells were treated with 10-NCP (inhibits Akt to pAkt conversion), it was observed that Caspr1 expression is downregulated in both cell lines (Figure 5C, D).

**Figure 5.**
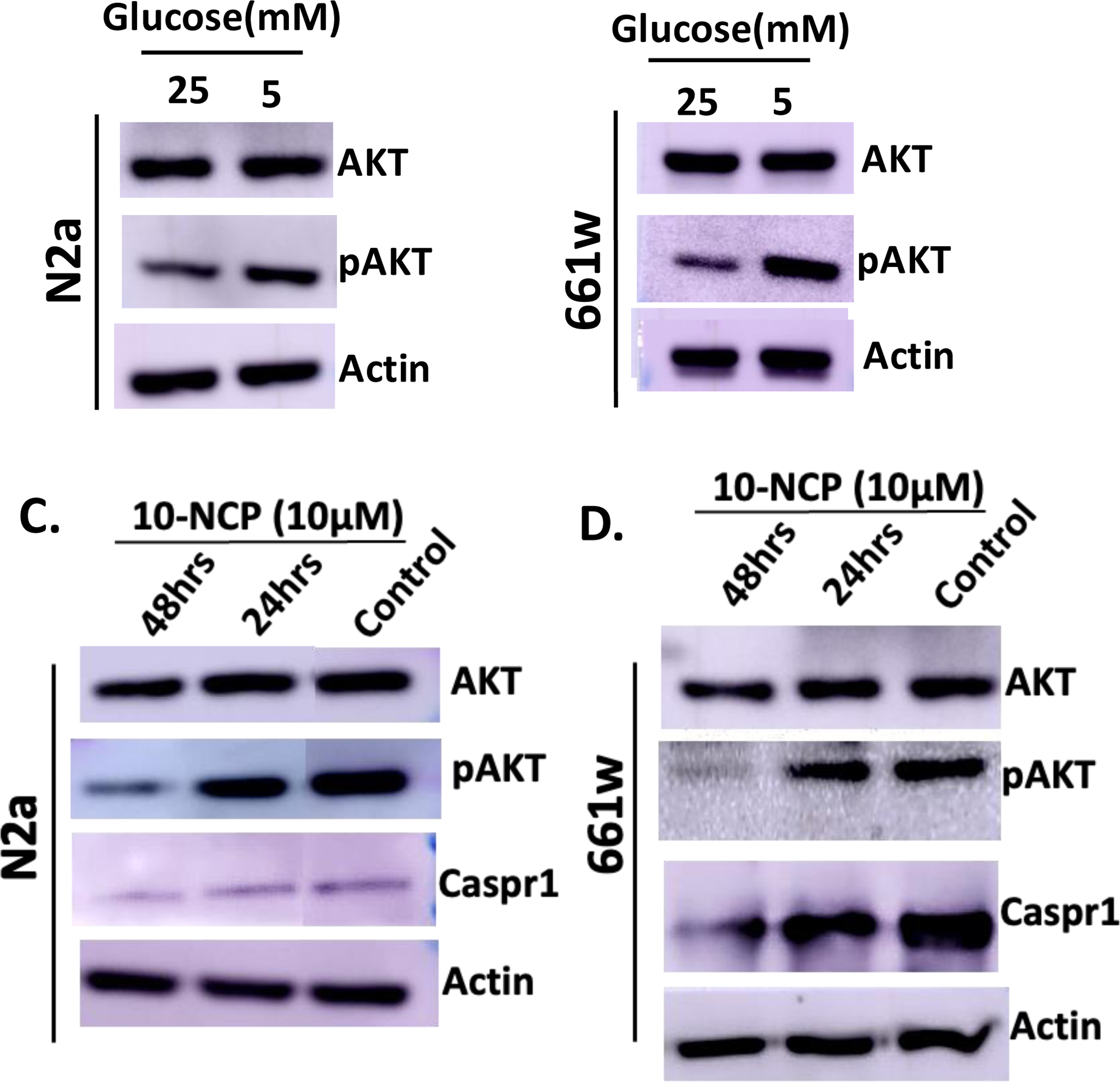
Hyperglycemia affects the Akt-dependent expression of Caspr-1. (A) and (B) Western blot analysis of Akt and pAkt protein expression in N2a and 661W cells grown in 5mM and 25mM glucose medium, respectively. (C) and (D) Western blot analysis of Akt and pAkt protein expression in N2a and 661W cells after treatment with Akt inhibitor (10-NCP) (10*μ*M), respectively.

### Insulin upregulates Caspr1 expression under hyperglycemic conditions

Insulin acts as a neuroprotective factor and helps in neuronal cell survival. Insulin activates the insulin receptor, followed by phosphatidylinositol 3-kinase (PI3K) and Akt, to promote cell survival. Emerging evidence indicates that insulin can enhance neuronal survival and aid recovery following trauma or in neurodegenerative diseases (Ravichandran et al., 2001; Apostolatos et al., 2012; Shaughness et al., 2020). Next, we examined the effect of insulin on Caspr1 expression as well as neurite length under hyperglycemic conditions. To do that, N2a and 661W cells were treated with different insulin concentrations (100nM, 200nM, 300nM) in the presence of 25mM glucose-containing media. Caspr1 mRNA expression was upregulated in 200nM concentration of insulin under hyperglycemic conditions (Supplementary Figure 4). Consistent with these results, we also noticed that Akt to phospho-Akt (pAkt) conversion and Caspr1 expression were upregulated in N2a and 661W cells (Figure 6A, B). Because Caspr1 expression was altered in the presence of insulin, neurite length was examined. Cells were grown in 25mM glucose-containing media with 200nM insulin for 48 hours, and a neurite outgrowth assay was performed. It was observed that cells had significantly shorter neurites in the presence of insulin than control; N2a cells (25mM: 47.7 ± 1.0 μm, 25mM+Insulin (200nM): 25.84 ± 0.67 μm (Figure 6D) and 661W cells (25mM: 48.6 ± 0.37 μm, 25mM+Insulin (200nM) 43.7 ± 1.76 μm) (Figure 6E).

**Figure 6.**
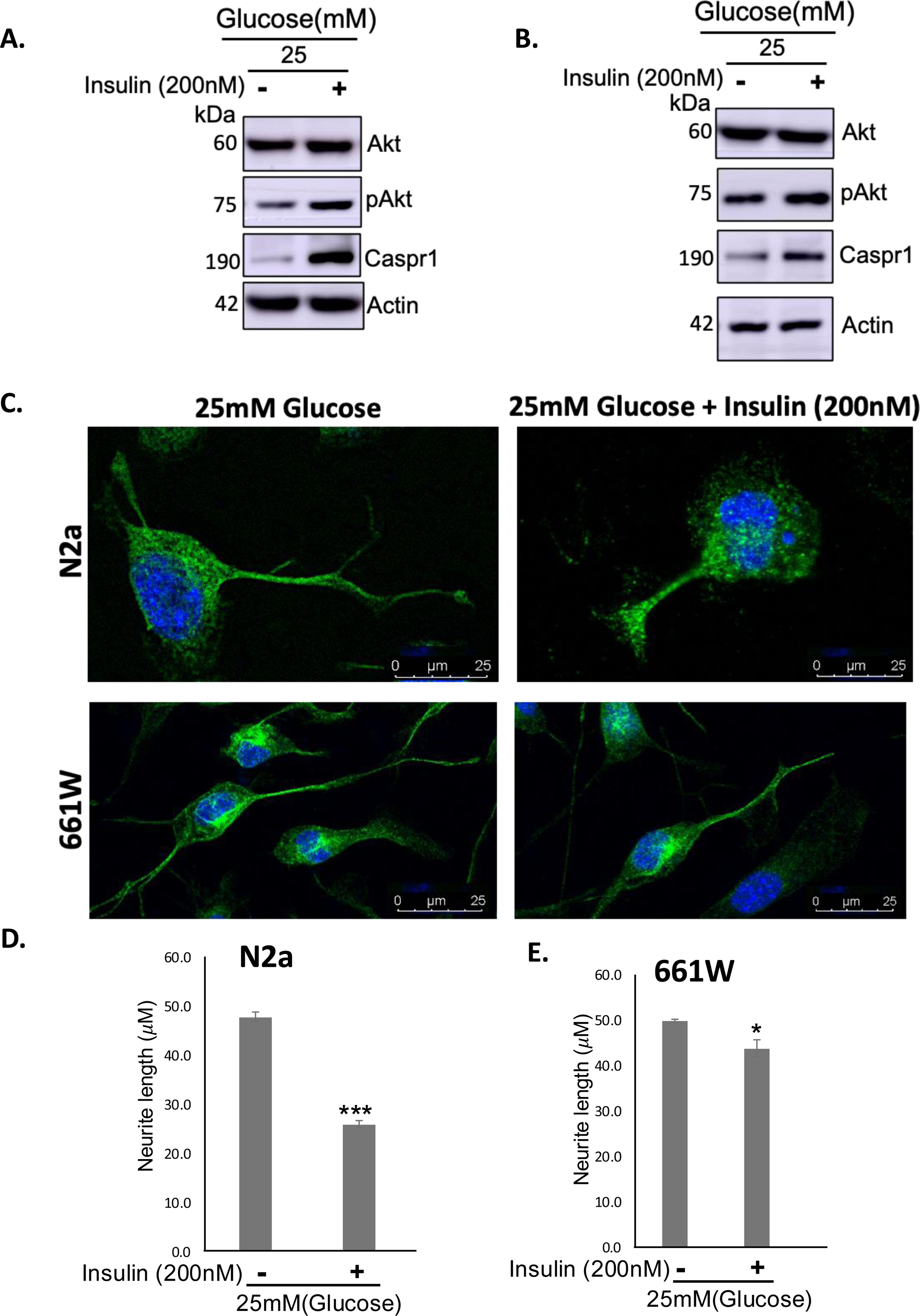
Insulin regulates Caspr1 expression and neurite length under hyperglycemic conditions. (A) and (B) Western blot analysis of Akt, pAkt and Caspr1 expression in N2a and 661W cells grown in 25mM glucose medium after the insulin (200nM) treatment, respectively. (C) Representative 40X *β*III tubulin (green) immunostained images showing neurite length in N2a and 661w cells cultured in 25mM glucose medium after the insulin (200nM) treatment. (D) and (E) Quantification of total neurite length shows a significant decrease in insulin-treated cells compared to control in N2a and 661w cells, respectively. N=3. *P<0.05, **P<0.01, ***P<0.001.

### Expression of Caspr1 in type-1 diabetic mice, high-fat diet mice and type II diabetic patients

Expression of Caspr1 and C/EBPα was also studied in the retina of C57BL/6 type-1 diabetic mice. To perform this experiment, three STZ-induced diabetic male C57BL/6 mice and three control male C57BL/6 mice were sacrificed. Eyeballs were removed carefully to make sections, and the retina was extracted to isolate RNA. The cross-section of the type 1 diabetic retina shows that Caspr1 expression is less in the outer layer of the retina in comparison with the control (Figure 6A). Further, we also checked the mRNA level of Caspr1 and C/EBPα in type-1 diabetic mice retina, and the result shows significant downregulation in the expression of Caspr1 and C/EBPα (Supplementary Figure 4A, B). Next, we examined the expression of Caspr1 in high-fat diet (HFD) induced obese mice. The immunostained cross-section of the HFD-fed mice retina showed less expression of Caspr1 in the inner layer and outer layer of the retina in comparison to the control mice (Figure 6B). We also observed similar results in type II diabetic patients (Figure 6C). From the above results, we concluded that Caspr1 expression gets downregulated in diabetic conditions.

## Discussion

Nearly half a billion people worldwide suffer from diabetes now, and the International Diabetes Federation has released new figures showing that by 2045, that number will rise to 629 million. The most frequent microvascular consequence of diabetes is retinopathy, which is also the main cause of blindness and vision impairment among working-age individuals (20–74 years old) (Yau et al., 2012). Risk factors for developing diabetic retinopathy (DR) include hyperglycemia, dyslipidemia, hypertension, and the duration of diabetes (Frank, 2014; Hammer and Busik, 2017). Although vascular abnormalities are the predominant clinical features of diabetic retinopathy (DR), early research studies have also documented neuropathic symptoms such as degeneration of the ganglion cell layers and inner nuclear layer, along with pathological changes in the nerve fiber layer. Long-term hyperglycemia in diabetes damages the blood-retinal barrier’s integrity, which results in altered permeability of the endothelial cells and pericytes lining retinal capillaries, altered autoregulation, and increased release of growth factors and cytokines (Chakravarthy and Devanathan, 2018). To explore the effect of hyperglycemia on retinal neurons, we first established a hyperglycemic model of adult retinal neurons and explored the intricate relationship between Caspr1, hyperglycemia, and neuritogenesis, revealing a significant role of CEBPα as a transcriptional regulator of Caspr1, which in turn influences neurite outgrowth (Figure 8). Caspr1 has been extensively studied in the brain for its involvement in controlling neurite outgrowth (Devanathan et al., 2010) and the nodal/paranodal domain organisation of myelinated axons (Peles and Salzer, 2000). However, Caspr1’s role in the retina, which is made up of non-myelinated neurones, remains unknown. Our findings demonstrate that hyperglycemic condition leads to the absence of Caspr1 (Figure 1A, B), and it is associated with increased neurite outgrowth in retinal neurons (Figure 2), suggesting that Caspr1 normally functions as an inhibitory regulator in this process not only in the brain but also in the retina. According to previous studies, the IL-1β-dependent mechanism may promote neurite extension in adult neurones (Saleh et al., 2013). Hyperglycemia has been linked to elevated levels of pro-inflammatory cytokines, including IL-1β and TNF-α, in the diabetic retina (Busik et al., 2008; Chakravarthy and Devanathan, 2018), which may impact neuritogenesis as seen in our work. This insight provides a critical understanding of the molecular mechanisms that govern neuritogenesis in the context of diabetes and metabolic disturbances.

The C/EBP family of transcription factors is sensitive to alterations in glucose homeostasis (Schroeder-Gloeckler et al., 2007). C/EBP α and β are abundant in CNS neurons and have been linked to neuronal development, survival, and neurogenesis (Ménard et al., 2002; Ramji and Foka, 2002). There are no reports about the importance of CEBPα in neurite outgrowth regulation, in particular, our study indicates the strong role of CEBPα in transcriptionally regulating the cell adhesion molecule Caspr1. Caspr1 is a huge protein (190 kDa) including glycosylation. Our results indicate an interesting regulatory mechanism for cell adhesion molecules. In addition, this is, to our knowledge, the first report where CEBPα regulates a transmembrane protein. When CEBPα was overexpressed in the cells, it upregulated the Caspr1 promoter activity and Caspr1 expression (Figure 4). CEBPα acts as a transcriptional enhancer for Caspr1 in normoglycemic conditions; however, under hyperglycemic conditions, CEBPα expression was reduced (Figure 4D, E), and thereby, Caspr1 expression was also downregulated(Figure 1A). This relationship highlights the potential of targeting the Caspr1-CEBPα pathway to influence neuronal development and regeneration, particularly in conditions characterized by metabolic dysregulation.

In diabetes, the most affected pathway is the insulin-Akt pathway. Under hyperglycemic conditions, pAkt expression gets downregulated, which affects cell survival (Ravichandran et al., 2001; Shaughness et al., 2020). In our study, when the cells were grown in hyperglycaemia and supplemented with insulin, pAkt expression increased, which led to upregulated Caspr1 expression (Figure 6A, B).

Caspr1 and Contactin1 usually form a complex at the paranodal junction of myelinated neurones in the CNS. The lack of Caspr1 causes Contactin1 to mislocalize and be excluded from the paranodes (Bhat et al., 2001), whereas the absence of Contactin1 prevents Caspr1 from reaching the axonal membrane (Boyle et al., 2001). Caspr1 is also related to neurodegenerative diseases such as multiple sclerosis and Alzheimer’s. In multiple sclerosis, Caspr1 expression gets downregulated, which leads to the diffusion of K+ channels, and the channels start overlapping with the paranodal region, which causes neurodegeneration (Coman et al., 2006; Zou et al., 2017). In our results, Caspr1 expression is downregulated in Type 1 diabetes mouse models and in diabetic patients (Figure 7A, C), suggesting that hyperglycemic conditions reduce Caspr1(Figure 1A) and affect neuronal morphology. Similarly, in high-fat diet mouse models, reduced Caspr1 expression aligns with the notion that metabolic disturbances can impact neuronal health by altering the expression of key regulatory proteins (Figure 7B).

**Figure 7.**
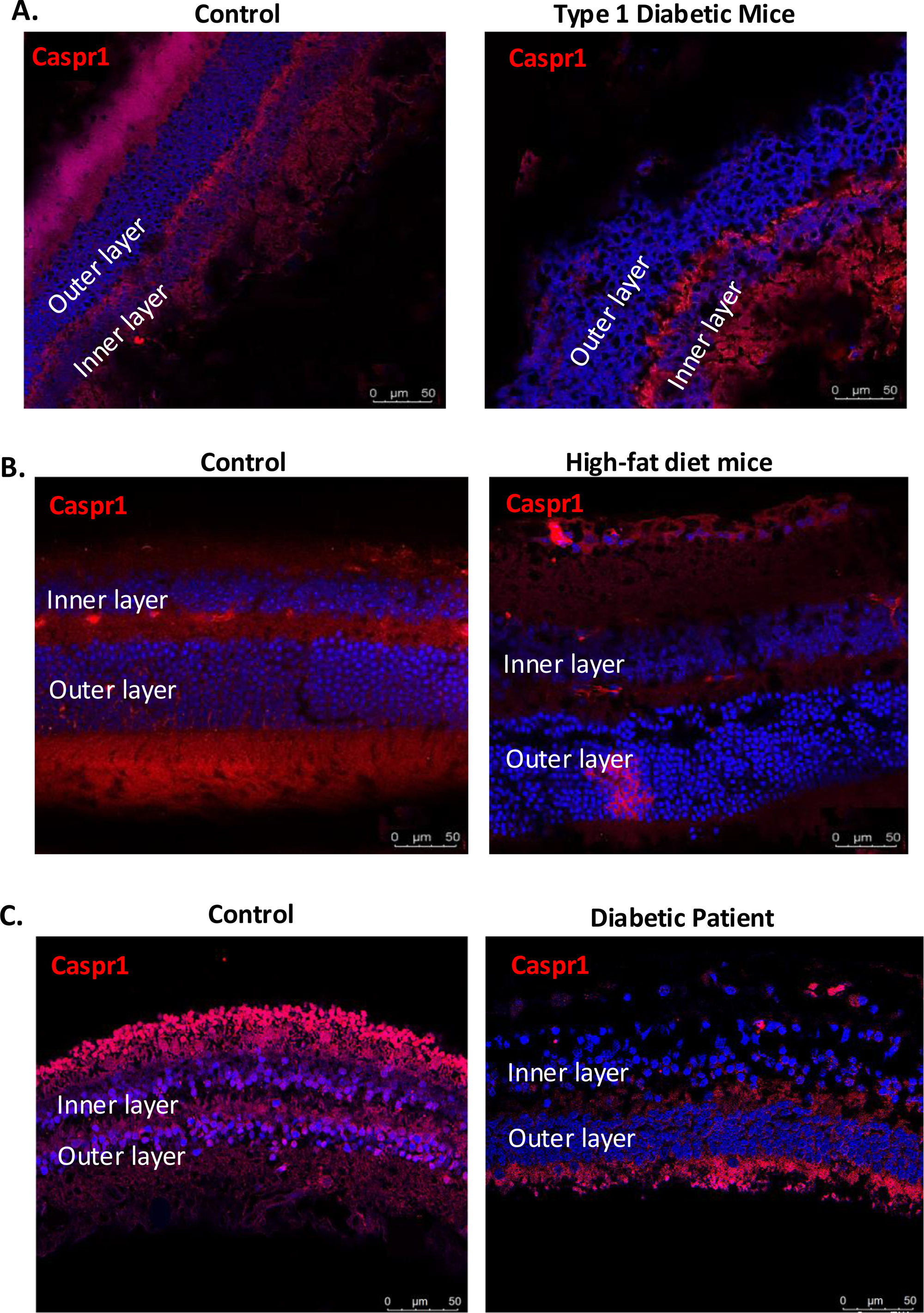
Expression of Caspr1 in diabetic retina. (A) Cross-section of type I diabetic mice retina stained with Caspr1 (Red) antibody. (B) Cross-section of high-fat diet mice retina stained with Caspr1 (Red) antibody. (C) Cross-section of the type II diabetic patient retina stained with Caspr1 (Red) antibody.

The enhanced neurite outgrowth observed in the absence of Caspr1 raises important questions regarding the balance between inhibitory and permissive signals in neuronal development. In the context of hyperglycemia, where oxidative stress and inflammation are prevalent, the absence of inhibitory factors like Caspr1 could facilitate compensatory growth responses in neurons. However, the long-term consequences of this increased outgrowth in a hyperglycemic environment warrant further investigation, as excessive neurite growth might disrupt normal neuronal networks and lead to functional impairments.

Additionally, the findings suggest that interventions aimed at modulating Caspr1 levels or its downstream pathways could have therapeutic potential. For instance, targeting Caspr1 expression or function may regulate neurite outgrowth in diabetic conditions, aiding neuronal recovery and plasticity. Conversely, understanding the pathways that lead to downregulating Caspr1 expression in hyperglycemic conditions could illuminate novel strategies for mitigating the adverse effects of diabetes on the nervous system (Figure 8).

**Figure 8.** Schematic diagram showing the regulation of Caspr1 expression and function under hyperglycemia. Schematic depiction of Caspr1 regulation under hyperglycemic conditions. High glucose concentration in the medium downregulates Caspr1 expression by affecting Akt signalling and C/EBPα. Hyperglycemia affects the expression of C/EBPα and causes a reduction in Caspr1 level in retinal neurons, responsible for longer and unregulated neurite outgrowth (left). Moreover, Caspr1 expression was upregulated by increased expression of pAkt in the presence of insulin under hyperglycemia, which leads to regulated neurite outgrowth (right).

In conclusion, our study highlights the pivotal role of Caspr1 in regulating neuritogenesis, particularly in the context of hyperglycemia and metabolic stress. The transcriptional regulation of Caspr1 by CEBPα presents a key mechanism by which Caspr1 modulates neurite outgrowth under hyperglycemia (Figure 8). These findings not only enhance our understanding of the molecular interactions governing neuritogenesis but also open new avenues for therapeutic interventions in diabetes-related neuronal dysfunction. Future research should further elucidate the signalling pathways involved and explore the therapeutic potential of manipulating Caspr1 and CEBPα in both preclinical and clinical settings.

## Conflict of Interest

The authors state that the study was done without any commercial or financial relationships that may be considered a possible conflict of interest.

## Author Contribution

SS conducted research, analysed data, and wrote the report. GS and SS conducted research and analysed data. VD conceived the study, provided reagents/analytic tools, and authored the manuscript. VD is the work’s guarantor; therefore had complete access to all of the study’s data. All authors accept responsibility for the data’s integrity and accuracy during analysis.

## Funding

This research was funded by IISER Tirupati and DST Department of Science and Technology, Govt of India. VD is a recipient of the Core Research Grant (CRG), Department of Science and Technology (DST). BSS is a recipient of ICGEB Trieste, NBRC core funds, SRG from SERB, and Grant IBRO, Paris. SS is a recipient of the DAAD fellowship.

## Acknowledgments

VD thanks IISER Tirupati for research support. SS thanks IISER Tirupati, UGC and DAAD for her research fellowship. We thank Dr. Pakala Suresh Babu’s excellent comments, which allowed us to expand on our study ideas.

---

**Video 1.** Live cell imaging of wild-type (upper) and Caspr1 knockout (down) cells. Caspr1 knockout cells were growing longer neurites compared to wild-type.

**Supplementary Figure 1.** Cloning of sgRNAs against Caspr1 gene into pU6-(BbsI)_CBh-Cas9-T2A-mCherry vector. (A) Cloning of sgRNA1 and sgRNA2 into pU6-(BbsI)_CBh-Cas9-T2A-mCherry vector by colony PCR amplification of 360bp from the construct. (B) After performing Gibson assembly, confirmation of sgRNA1 and sgRNA2 into pU6-(BbsI)_CBh-Cas9-T2A-mCherry vector by colony PCR amplification of 500bp from the construct. (C) Representative image of N2a cells transfected with sgRNAs containing pU6-(BbsI)_CBh-Cas9-T2A-mCherry vector expressing mCherry.

**Supplementary Figure 2.** Cloning of mouse Caspr1 promoter (-1 to -2000bp) into pGL2 basic vector. (A) PCR amplification of the mouse Caspr1 promoter from the genomic DNA isolated from 661W cells. (B) Restriction digestion of the pGL2 vector with *SmaI* and *XhoI* enzymes.

**Supplementary Figure 3.** PCR amplification of C/EBPα FL from the cDNA isolated from 661W cells.

**Supplementary Figure 4.** (A, B) qPCR analysis of Caspr1 expression in different concentrations of insulin in N2a and 661W cells cultured in 25mM glucose-containing media, respectively. β-Actin was used for normalisation. The degree of significance is represented as follows: ***P<0.001, **P<0.01 and *P<0.05.

**Supplementary Figure 5.** (A, B) qPCR analysis of Caspr1 and C/EBPα mRNA expression in Control and diabetic mice, respectively. β-Actin was used for normalization. The degree of significance is represented as follows: ***P<0.001, **P<0.01 and *P<0.05.

## Notes

### Competing Interest Statement

The authors have declared no competing interest.

## Reference

Apostolatos, A., Song, S., Acosta, S., Peart, M., Watson, J.E., Bickford, P., Cooper, D.R., Patel, N.A., 2012. Insulin Promotes Neuronal Survival via the Alternatively Spliced Protein Kinase CδII Isoform. J. Biol. Chem. 287, 9299. 10.1074/jbc.M111.313080

Bhat, M.A., Rios, J.C., Lu, Y., Garcia-Fresco, G.P., Ching, W., St Martin, M., Li, J., Einheber, S., Chesler, M., Rosenbluth, J., Salzer, J.L., Bellen, H.J., 2001. Axon-glia interactions and the domain organization of myelinated axons requires neurexin IV/Caspr/Paranodin. Neuron 30, 369–383. 10.1016/s0896-6273(01)00294-x

Boyle, M.E., Berglund, E.O., Murai, K.K., Weber, L., Peles, E., Ranscht, B., 2001. Contactin orchestrates assembly of the septate-like junctions at the paranode in myelinated peripheral nerve. Neuron 30, 385–397. 10.1016/s0896-6273(01)00296-3

Bromberg, K.D., Iyengar, R., He, J.C., 2008. Regulation of neurite outgrowth by Gi/o signaling pathways. Front. Biosci. J. Virtual Libr. 13, 4544–4557.

Busik, J.V., Mohr, S., Grant, M.B., 2008. Hyperglycemia-induced reactive oxygen species toxicity to endothelial cells is dependent on paracrine mediators. Diabetes 57, 1952–1965. 10.2337/db07-1520

Chakravarthy, H., Devanathan, V., 2018. Molecular Mechanisms Mediating Diabetic Retinal Neurodegeneration: Potential Research Avenues and Therapeutic Targets. J. Mol. Neurosci. MN 66, 445–461. 10.1007/s12031-018-1188-x

Coman, I., Aigrot, M.S., Seilhean, D., Reynolds, R., Girault, J.A., Zalc, B., Lubetzki, C., 2006. Nodal, paranodal and juxtaparanodal axonal proteins during demyelination and remyelination in multiple sclerosis. Brain J. Neurol. 129, 3186–3195. 10.1093/brain/awl144

Dadon-Nachum, M., Melamed, E., Offen, D., 2011. The “dying-back” phenomenon of motor neurons in ALS. J. Mol. Neurosci. MN 43, 470–477. 10.1007/s12031-010-9467-1

Devanathan, V., Jakovcevski, I., Santuccione, A., Li, S., Lee, H.J., Peles, E., Leshchyns’ka, I., Sytnyk, V., Schachner, M., 2010. Cellular Form of Prion Protein Inhibits Reelin-Mediated Shedding of Caspr from the Neuronal Cell Surface to Potentiate Caspr-Mediated Inhibition of Neurite Outgrowth. J. Neurosci. 30, 9292–9305. 10.1523/JNEUROSCI.5657-09.2010

Elliott, R.C., Miles, M.F., Lowenstein, D.H., 2003. Overlapping microarray profiles of dentate gyrus gene expression during development- and epilepsy-associated neurogenesis and axon outgrowth. J. Neurosci. Off. J. Soc. Neurosci. 23, 2218–2227. 10.1523/JNEUROSCI.23-06-02218.2003

Fan, L., Xu, D., Wang, W., Yan, K., Wu, H., Yao, X., Xu, R., Liu, C., Ma, Q., 2013. Caspr interaction with Amyloid Precursor Protein reduces amyloid-β generation in vitro. Neurosci. Lett. 548, 255–260. 10.1016/j.neulet.2013.05.055

Frank, R.N., 2014. Systemic Therapies for Diabetic Retinopathy: The Accord Eye Study. Ophthalmology 121, 2295–2296. 10.1016/j.ophtha.2014.08.019

Govek, E.-E., Newey, S.E., Aelst, L.V., 2005. The role of the Rho GTPases in neuronal development. Genes Dev. 19, 1–49. 10.1101/gad.1256405

Hammer, S.S., Busik, J.V., 2017. The role of dyslipidemia in diabetic retinopathy. Vision Res., Diabetic Retinopathy - an Overview 139, 228–236. 10.1016/j.visres.2017.04.010

Howell, O.W., Palser, A., Polito, A., Melrose, S., Zonta, B., Scheiermann, C., Vora, A.J., Brophy, P.J., Reynolds, R., 2006. Disruption of neurofascin localization reveals early changes preceding demyelination and remyelination in multiple sclerosis. Brain J. Neurol. 129, 3173–3185. 10.1093/brain/awl290

Huang, E.J., Reichardt, L.F., 2001. Neurotrophins: roles in neuronal development and function. Annu. Rev. Neurosci. 24, 677–736. 10.1146/annurev.neuro.24.1.677

Hur, J.-Y., Teranishi, Y., Kihara, T., Yamamoto, N.G., Inoue, M., Hosia, W., Hashimoto, M., Winblad, B., Frykman, S., Tjernberg, L.O., 2012. Identification of Novel γ-Secretase-associated Proteins in Detergent-resistant Membranes from Brain. J. Biol. Chem. 287, 11991–12005. 10.1074/jbc.M111.246074

Jones, B.W., Watt, C.B., Frederick, J.M., Baehr, W., Chen, C.-K., Levine, E.M., Milam, A.H., Lavail, M.M., Marc, R.E., 2003. Retinal remodeling triggered by photoreceptor degenerations. J. Comp. Neurol. 464, 1–16. 10.1002/cne.10703

Lin, Y., Jones, B.W., Liu, A., Tucker, J.F., Rapp, K., Luo, L., Baehr, W., Bernstein, P.S., Watt, C.B., Yang, J.-H., Shaw, M.V., Marc, R.E., 2012. Retinoid receptors trigger neuritogenesis in retinal degenerations. FASEB J. 26, 81–92. 10.1096/fj.11-192914

Ling-Sing Seow, S., Naidu, M., David, P., Wong, K.-H., Sabaratnam, V., 2013. Potentiation of neuritogenic activity of medicinal mushrooms in rat pheochromocytoma cells. BMC Complement. Altern. Med. 13, 157. 10.1186/1472-6882-13-157

Marc, R.E., Jones, B.W., 2003. Retinal remodeling in inherited photoreceptor degenerations. Mol. Neurobiol. 28, 139–147. 10.1385/MN:28:2:139

Marc, R.E., Jones, B.W., Anderson, J.R., Kinard, K., Marshak, D.W., Wilson, J.H., Wensel, T., Lucas, R.J., 2007. Neural reprogramming in retinal degeneration. Invest. Ophthalmol. Vis. Sci. 48, 3364–3371. 10.1167/iovs.07-0032

Marc, R.E., Jones, B.W., Watt, C.B., Strettoi, E., 2003. Neural remodeling in retinal degeneration. Prog. Retin. Eye Res. 22, 607–655. 10.1016/s1350-9462(03)00039-9

Ménard, C., Hein, P., Paquin, A., Savelson, A., Yang, X.M., Lederfein, D., Barnabé-Heider, F., Mir, A.A., Sterneck, E., Peterson, A.C., Johnson, P.F., Vinson, C., Miller, F.D., 2002. An essential role for a MEK-C/EBP pathway during growth factor-regulated cortical neurogenesis. Neuron 36, 597–610. 10.1016/s0896-6273(02)01026-7

More, S.V., Koppula, S., Kim, I.-S., Kumar, H., Kim, B.-W., Choi, D.-K., 2012. The role of bioactive compounds on the promotion of neurite outgrowth. Mol. Basel Switz. 17, 6728–6753. 10.3390/molecules17066728

Peles, E., Salzer, J.L., 2000. Molecular domains of myelinated axons. Curr. Opin. Neurobiol. 10, 558–565. 10.1016/s0959-4388(00)00122-7

Rai, S.N., Dilnashin, H., Birla, H., Singh, S.S., Zahra, W., Rathore, A.S., Singh, B.K., Singh, S.P., 2019. The Role of PI3K/Akt and ERK in Neurodegenerative Disorders. Neurotox. Res. 35, 775–795. 10.1007/s12640-019-0003-y

Ramji, D.P., Foka, P., 2002. CCAAT/enhancer-binding proteins: structure, function and regulation. Biochem. J. 365, 561–575. 10.1042/BJ20020508

Ravichandran, L.V., Esposito, D.L., Chen, J., Quon, M.J., 2001. Protein kinase C-zeta phosphorylates insulin receptor substrate-1 and impairs its ability to activate phosphatidylinositol 3-kinase in response to insulin. J. Biol. Chem. 276, 3543–3549. 10.1074/jbc.M007231200

Read, D.E., Gorman, A.M., 2009. Involvement of Akt in neurite outgrowth. Cell. Mol. Life Sci. 66, 2975–2984. 10.1007/s00018-009-0057-8

Saleh, A., Chowdhury, S.K.R., Smith, D.R., Balakrishnan, S., Tessler, L., Schartner, E., Bilodeau, A., Van Der Ploeg, R., Fernyhough, P., 2013. Diabetes impairs an interleukin-1β-dependent pathway that enhances neurite outgrowth through JAK/STAT3 modulation of mitochondrial bioenergetics in adult sensory neurons. Mol. Brain 6, 45. 10.1186/1756-6606-6-45

Sayyad, Z., Sirohi, K., Radha, V., Swarup, G., 2017. 661W is a retinal ganglion precursor-like cell line in which glaucoma-associated optineurin mutants induce cell death selectively. Sci. Rep. 7, 16855. 10.1038/s41598-017-17241-0

Scheibel, A.B., Tomiyasu, U., 1978. Dendritic sprouting in Alzheimer’s presenile dementia. Exp. Neurol. 60, 1–8. 10.1016/0014-4886(78)90164-4

Schroeder-Gloeckler, J.M., Rahman, S.M., Janssen, R.C., Qiao, L., Shao, J., Roper, M., Fischer, S.J., Lowe, E., Orlicky, D.J., McManaman, J.L., Palmer, C., Gitomer, W.L., Huang, W., O’Doherty, R.M., Becker, T.C., Klemm, D.J., Jensen, D.R., Pulawa, L.K., Eckel, R.H., Friedman, J.E., 2007. CCAAT/enhancer-binding protein beta deletion reduces adiposity, hepatic steatosis, and diabetes in Lepr(db/db) mice. J. Biol. Chem. 282, 15717–15729. 10.1074/jbc.M701329200

Sharma, S., Chakravarthy, H., Suresh, G., Devanathan, V., 2019. Adult Goat Retinal Neuronal Culture: Applications in Modeling Hyperglycemia. Front. Neurosci. 13.

Shaughness, M., Acs, D., Brabazon, F., Hockenbury, N., Byrnes, K.R., 2020. Role of Insulin in Neurotrauma and Neurodegeneration: A Review. Front. Neurosci. 14. 10.3389/fnins.2020.547175

Tang, S.-Y., Liu, D.-X., Li, Y., Wang, K.-J., Wang, X.-F., Su, Z.-K., Fang, W.-G., Qin, X.-X., Wei, J.-Y., Zhao, W.-D., Chen, Y.-H., 2020. Caspr1 Facilitates sAPPα Production by Regulating α-Secretase ADAM9 in Brain Endothelial Cells. Front. Mol. Neurosci. 13.

Varma, S., Lal, B.K., Zheng, R., Breslin, J.W., Saito, S., Pappas, P.J., Hobson, R.W., Durán, W.N., 2005. Hyperglycemia alters PI3k and Akt signaling and leads to endothelial cell proliferative dysfunction. Am. J. Physiol. Heart Circ. Physiol. 289, H1744–H1751. 10.1152/ajpheart.01088.2004

Yau, J.W.Y., Rogers, S.L., Kawasaki, R., Lamoureux, E.L., Kowalski, J.W., Bek, T., Chen, S.-J., Dekker, J.M., Fletcher, A., Grauslund, J., Haffner, S., Hamman, R.F., Ikram, M.K., Kayama, T., Klein, B.E.K., Klein, R., Krishnaiah, S., Mayurasakorn, K., O’Hare, J.P., Orchard, T.J., Porta, M., Rema, M., Roy, M.S., Sharma, T., Shaw, J., Taylor, H., Tielsch, J.M., Varma, R., Wang, J.J., Wang, N., West, S., Xu, L., Yasuda, M., Zhang, X., Mitchell, P., Wong, T.Y., for the Meta-Analysis for Eye Disease (META-EYE) Study Group, 2012. Global Prevalence and Major Risk Factors of Diabetic Retinopathy. Diabetes Care 35, 556–564. 10.2337/dc11-1909

Zou, Y., Zhang, W., Liu, H., Li, X., Zhang, X., Ma, X., Sun, Y., Jiang, S., Ma, Q., Xu, D., 2017. Structure and function of the contactin-associated protein family in myelinated axons and their relationship with nerve diseases. Neural Regen. Res. 12, 1551–1558. 10.4103/1673-5374.215268

