## Supplemental Fig 1 for "Hyperglycemia transcriptionally regulates the paranodal protein (Caspr1) in retinal neurons and modulates neurite extension"

### Cloning of sgRNAs against Caspr1 gene into pU6-(BbsI)_CBh-Cas9-T2A-mCherry vector

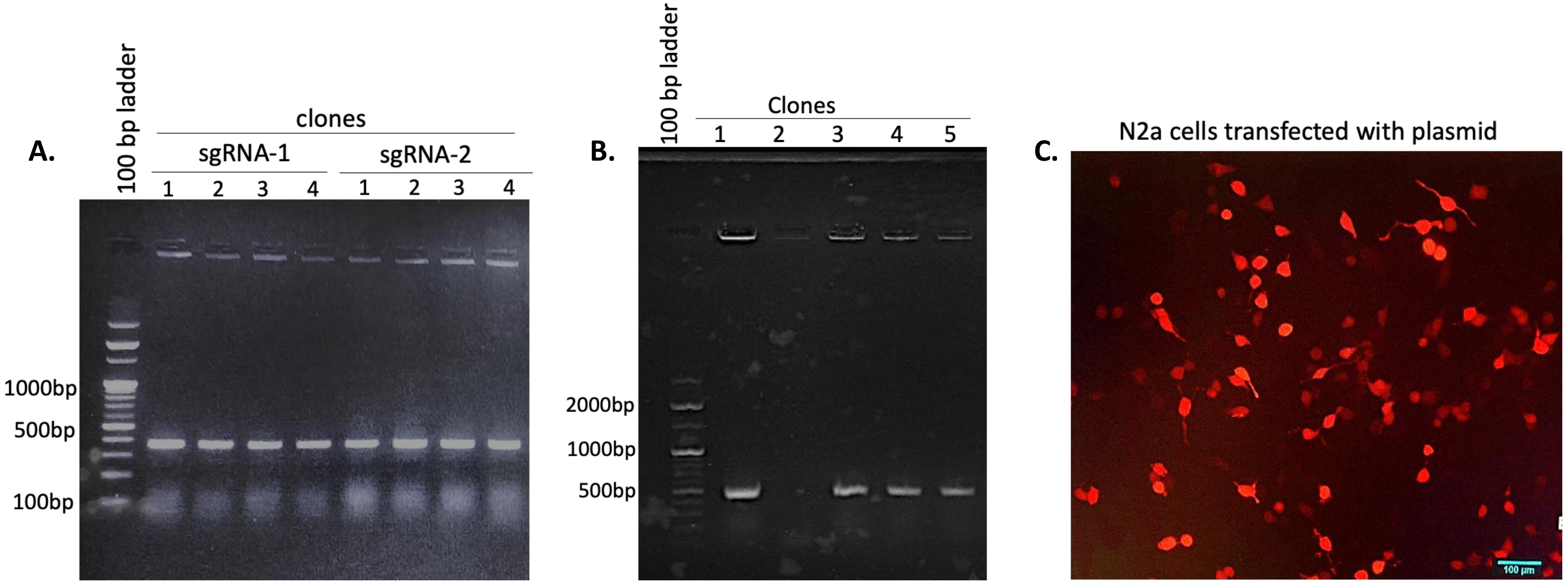


**Figure 1- Cloning of sgRNAs against Caspr1 gene into pU6-(BbsI)_CBh-Cas9-T2A-mCherry vector.** (A) Cloning of sgRNA1 and sgRNA2 into pU6-(BbsI)_CBh-Cas9-T2A-mCherry vector by colony PCR amplification of 360bp from the construct. (B) After performing Gibson assembly, confirmation of sgRNA1 and sgRNA2 into pU6-(BbsI)_CBh-Cas9-T2A-mCherry vector by colony PCR amplification of 500bp from the construct. (C) Representative image of N2a cells transfected with sgRNAs containing pU6-(BbsI)_CBh-Cas9-T2A-mCherry vector expressing mCherry.
