## Supplemental Fig 2 for "Hyperglycemia transcriptionally regulates the paranodal protein (Caspr1) in retinal neurons and modulates neurite extension"

### Cloning of mouse Caspr1 promoter (-1 to -2000bp) into pGL2 basic vector

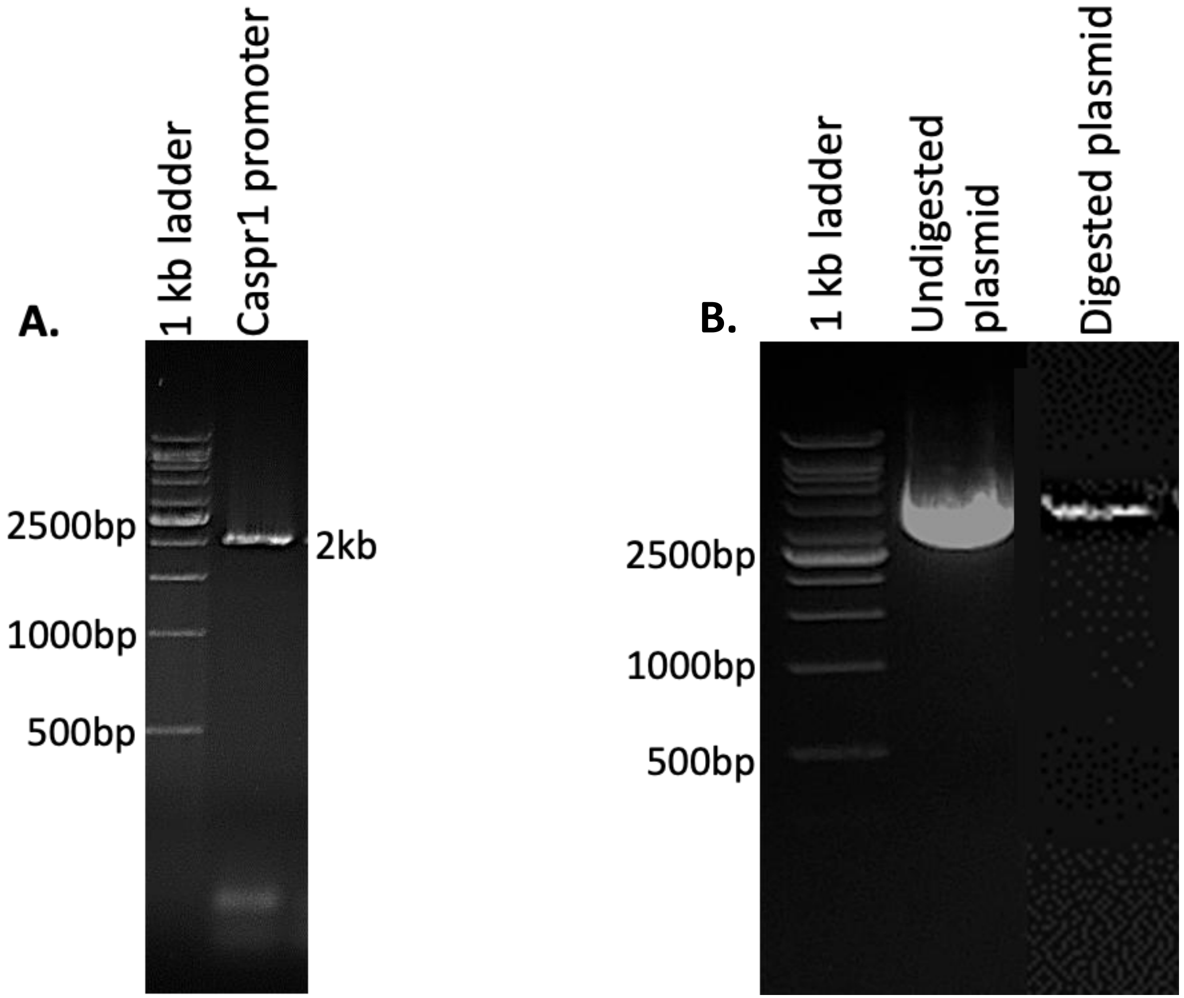


**Cloning of mouse Caspr1 promoter (-1 to -2000bp) into pGL2 basic vector. (A)** PCR amplification of mouse Caspr1 promoter from the genomic DNA isolated from 661W cells. **(B)** Restriction digestion of pGL2 vector with *SmaI* and *XhoI* enzymes.
