## Supplemental Fig 3 for "Hyperglycemia transcriptionally regulates the paranodal protein (Caspr1) in retinal neurons and modulates neurite extension"

### Amplification of  C/EBPα Full length

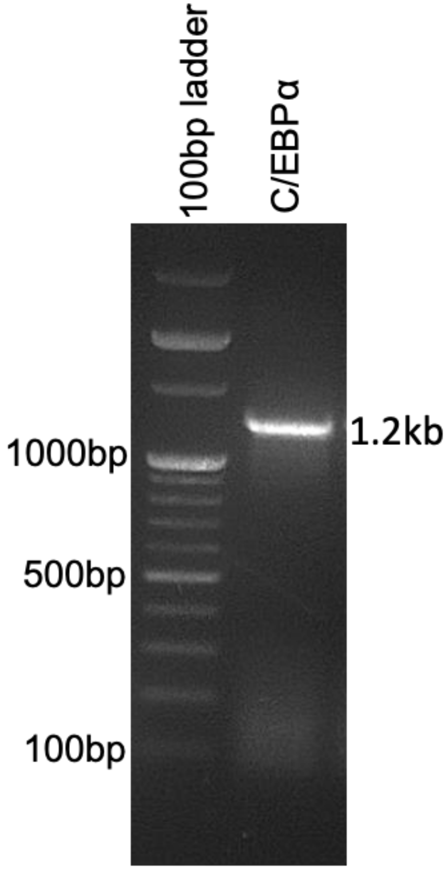


PCR amplification of C/EBPα FL from the cDNA isolated from 661W cells.
