## Supplemental Fig 4 for "Hyperglycemia transcriptionally regulates the paranodal protein (Caspr1) in retinal neurons and modulates neurite extension"

### Caspr1 expression in different insulin concentrations under hyperglycemia

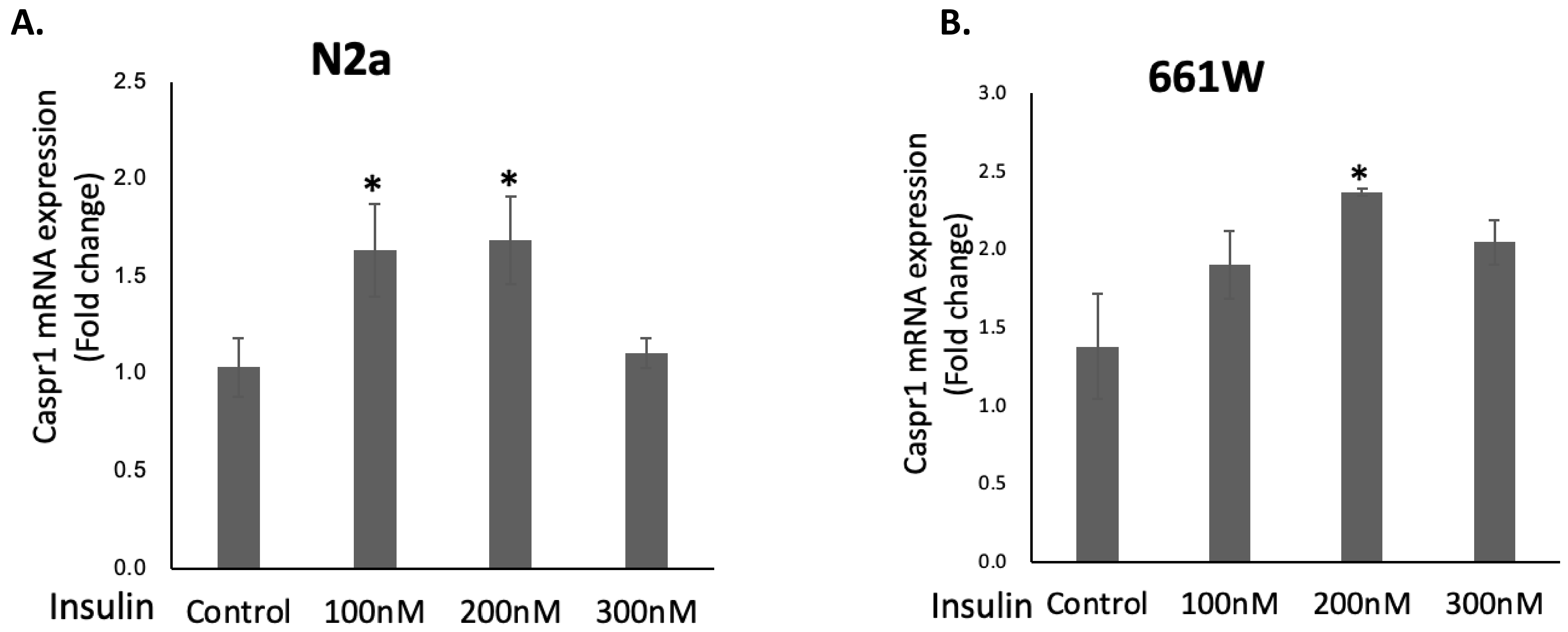


(A, B) qPCR analysis of Caspr1 expression in different concentrations of insulin in N2a and 661W cells cultured in 25mM glucose-containing media, respectively. β-Actin was used for normalisation. The degree of significance is represented as follows: ***$P<0.001$, **$P<0.01$ and *$P<0.05$.
