## Supplemental Fig 5 for "Hyperglycemia transcriptionally regulates the paranodal protein (Caspr1) in retinal neurons and modulates neurite extension"

### Caspr1 and C/EBPα expression in type 1 diabetic mice

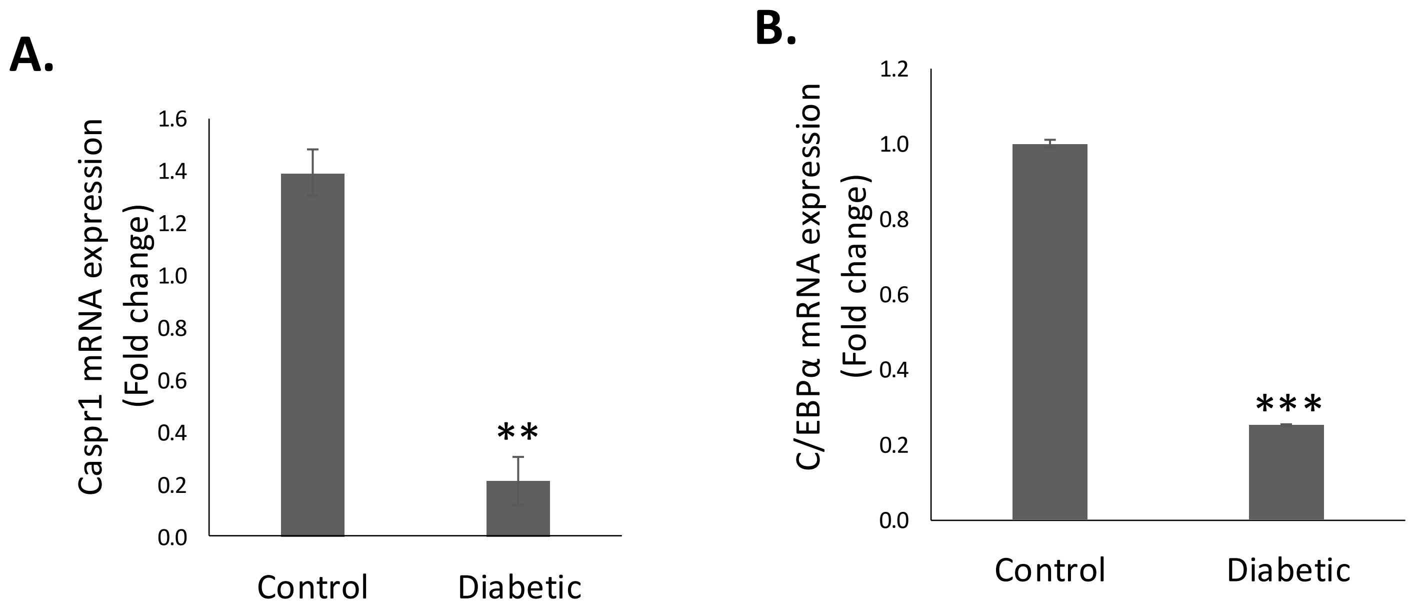


(A, B) qPCR analysis of Caspr1 and C/EBPα mRNA expression in Control and diabetic mice, respectively. β-Actin was used for normalization. The degree of significance is represented as follows: ***$P<0.001$, **$P<0.01$ and *$P<0.05$.
